# Disruption of a *Plasmodium falciparum* patatin-like phospholipase delays male gametocyte exflagellation

**DOI:** 10.1101/2023.04.28.538693

**Authors:** Emma Pietsch, Korbinian Niedermüller, Tim-Wolf Gilberger, Paul-Christian Burda

## Abstract

An essential process in transmission of the malaria parasite to the *Anopheles* vector is the conversion of mature gametocytes into gametes within the mosquito gut, where they egress from the red blood cell (RBC). During egress, male gametocytes undergo exflagellation, leading to the formation of eight haploid motile microgametes, while female gametes retain their spherical shape. Gametocyte egress depends on sequential disruption of the parasitophorous vacuole membrane and the host cell membrane. In other life cycle stages of the malaria parasite, phospholipases have been implicated in membrane disruption processes during egress, however their importance for gametocyte egress is relatively unknown. Here, we performed comprehensive functional analyses of six putative phospholipases for their role during development and egress of *Plasmodium falciparum* gametocytes. We localize two of them, the prodrug activation and resistance esterase (PF3D7_0709700) and the lysophospholipase 1 (PF3D7_1476700), to the parasite plasma membrane. Subsequently, we show that disruption of most of the studied phospholipase genes does neither affect gametocyte development nor egress. The exception is the putative patatin-like phospholipase PF3D7_0924000, whose gene deletion leads to a delay in male gametocyte exflagellation, indicating an important, albeit not essential, role of this enzyme in male gametogenesis.

## INTRODUCTION

Egress from gametocytes represents an essential process for the transmission of the human malaria parasite *Plasmodium falciparum* to its mosquito vector. It is induced by ingestion of mature gametocytes by female *Anopheles* mosquitoes during a blood meal. Within minutes after ingestion, gametocytes become activated in the mosquito midgut by a drop in temperature, a rise in pH, and the presence of the mosquito-derived molecule xanthurenic acid (Bennink *et al*., 2016). Following their activation, they undergo gametogenesis, whereby each female gametocyte produces a single immotile macrogamete, whereas a male gametocyte produces eight flagella-like microgametes in a process called exflagellation. After their release from host red blood cells (RBCs), macrogemetes and microgametes fuse to form the zygote, which later develops into the motile ookinete (Kuehn and Pradel, 2010).

During their intracellular development, gametocytes are surrounded by a parasitophorous vacuole membrane (PVM), which needs to be ruptured before and in addition to the RBC membrane for efficient release of gametes. Membrane rupture is facilitated by the exocytosis of specialized secretory vesicles of the parasites. These include the osmiophilic bodies that release a variety of egress-related proteins into the parasitophorous vacuole lumen (Flieger *et al*., 2018). In addition, other vesicles are released during egress than contain the perforin-like protein PPLP2, which is necessary for erythrocyte lysis (Deligianni *et al*., 2013; Wirth *et al*., 2014).

In other life cycle stages of malaria parasites, membrane rupture has been shown to be partially mediated by a secreted phospholipase with a lecithin:cholesterol acyltransferase-like domain. Rodent malaria *P. berghei* parasites missing this enzyme were defective in liver stage egress due to impaired PVM rupture (Burda *et al*., 2015) and conditional knockout of the same enzyme in *P. falciparum* negatively affected parasite release of asexual blood stage parasites from RBCs (Ramaprasad *et al*., 2023). The importance of phospholipases for membrane disruption processes during gametocyte egress is relatively unknown. Until now, only the patatin-like phospholipase 1 (PNPLA1) PF3D7_0209100 has a known function in this regard. Its conditional knockout reduced efficiency of gametocyte egress possibly by interfering with the release of egress-associated vesicles (Singh *et al*., 2019).

In this study, we performed a comprehensive functional analysis of six putative phospholipases for their role during development and egress of *P. falciparum* gametocytes. We show that five of the six phospholipases are dispensable for these processes but reveal that parasites lacking the putative PNPLA PF3D7_0924000 show a significant delay in efficient exflagellation of male gametocytes, suggesting an important but not essential function of this enzyme in male gametogenesis.

## RESULTS

### *P. falciparum* expresses at least 15 putative phospholipases during gametocyte development

The genome of *P. falciparum* includes 26 genes that encode for proteins containing putative lipase/phospholipase-related domains (PlasmoDB.org, (Aurrecoechea *et al*., 2009)). For 15 of these, there exists mass spectrometric evidence for expression in gametocytes (Florens *et al*., 2002; Silvestrini *et al*., 2010; Lasonder *et al*., 2016) (Figure 1), indicating a potential function in this parasite stage. Six of these gametocyte-expressed enzymes have a predicted signal peptide or contain transmembrane domains, suggesting they might be targeted into the secretory pathway and could be involved in membrane modification and/or disruption during the egress process (Figure 1). Given our previous results that disruption of the PF3D7_0908000 homologue in *P. berghei* did not result in a phenotype throughout the parasite life cycle (Burda *et al*., 2017) and the fact that the putative PNPLA2 (PF3D7_1358000) is important for mitochondrial function (Burda *et al*., 2021), we focused on the remaining four proteins (PF3D7_0731800, PF3D7_0924000, PF3D7_1411900, PF3D7_1412000) containing a predicted signal peptide or transmembrane domains for functional characterization in gametocytes. Additionally, we included the putative phospholipases PF3D7_1476700 and PF3D7_0709700 into our assays, as both show peak expression in stage V gametocytes based on RNAseq analysis (López-Barragán *et al*., 2011) (Figure 1).

**FIGURE 1.**
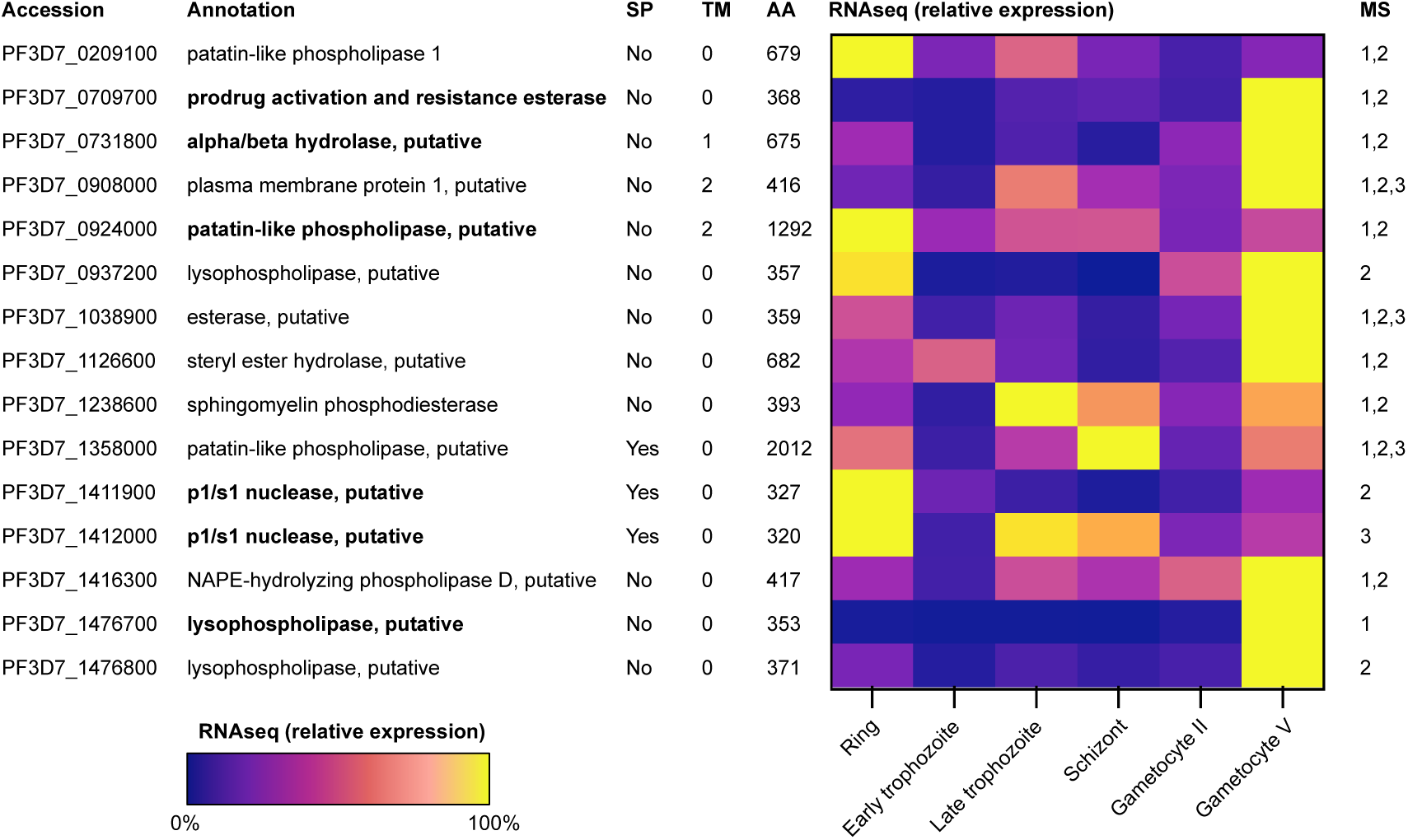
Putative phospholipases of *P. falciparum* that are expressed during gametocyte development based on mass spectrometry (MS). SP, signal peptide; TM, number of transmembrane domains; AA, length in amino acids. RNAseq data is based on (López-Barragán *et al*., 2011). Expression data derived from unique and non-unique reads was combined and normalized for each gene to the respective parasite stage showing highest expression, which was always set to 100%. MS data is based on: 1) (Lasonder *et al*., 2016); 2) (Silvestrini *et al*., 2010); 3) (Florens *et al*., 2002). Putative phospholipases analyzed in this study are shown in bold.

### The six selected putative phospholipases show various localization patterns in blood stage parasites

To study protein localization of the six selected putative phospholipases, we performed endogenous C-terminal tagging with mScarlet using the selection-linked integration (SLI) system (Birnbaum *et al*., 2017) in NF54/iGP2 parasites (Boltryk *et al*., 2021). This parasite line enables the production of high numbers of synchronous gametocytes and is transmission-competent, which means that it has maintained its ability to form gametes. Gametocyte commitment in this line is induced by conditional overexpression of the sexual commitment factor GDV1 (Boltryk *et al*., 2021). Correct integration of the respective constructs was verified by PCR (Figure S1) and localization analysis was performed by live-cell microscopy.

The prodrug activation and resistance esterase (PARE, PF3D7_0709700) showed a peripheral localization throughout asexual and sexual blood stage development that was also visible in free merozoites, suggesting that PARE localizes to the parasite plasma membrane (PPM) (Figure 2A). The lysophospholipase 4 (LPL4, PF3D7_0731800) (Asad *et al*., 2021) was not visible in ring and trophozoite stage parasites, while schizont stages displayed a weak punctuate pattern of LPL4-mScarlet in the cytoplasm and some signal in the food vacuole. During gametocyte development, LPL4-mScarlet localized mainly to the food vacuole. In addition, a peripheral signal was visible in some gametocytes (Figure 2B). Importantly, a similar fluorescence was absent when parental NF54/iGP2 wildtype parasites were imaged using the same settings, excluding that the observed signal is derived from unspecific background fluorescence (Figure S2). The putative PNPLA PF3D7_0924000, from here on referred to as PNPLA3, since PNPLA1 and PNPLA2 have already been described (Singh *et al*., 2019; Flammersfeld *et al*., 2020; Burda *et al*., 2021), could not be detected in asexual stages except for some autofluorescence of the hemozoin within the food vacuole. In gametocytes, a vesicular and diffuse cytoplasmic signal of PNPLA3-mScarlet was observed (Figure 3A). The putative phospholipase PF3D7_1411900, here termed phospholipase 39 (PL39) due to its predicted molecular weight of 39 kDa, localized to the food vacuole in all asexual and sexual blood stage parasites (Figure 3B). Interestingly, another putative phospholipase (PF3D7_1412000) is expressed from the neighboring gene locus. It has a predicted molecular weight of 38 kDa and is hence referred to as phospholipase 38 (PL38). In contrast to PL39, PL38-mScarlet displayed a punctuate localization pattern in schizonts and free merozoites, possibly suggesting a localization to the apical organelles of the parasite. In all other intraerythrocytic asexual stages and gametocytes no signal or only autofluorescence of the hemozoin could be observed, supporting a potential schizont specific function of PL38 (Figure 4A).

**FIGURE 2.**
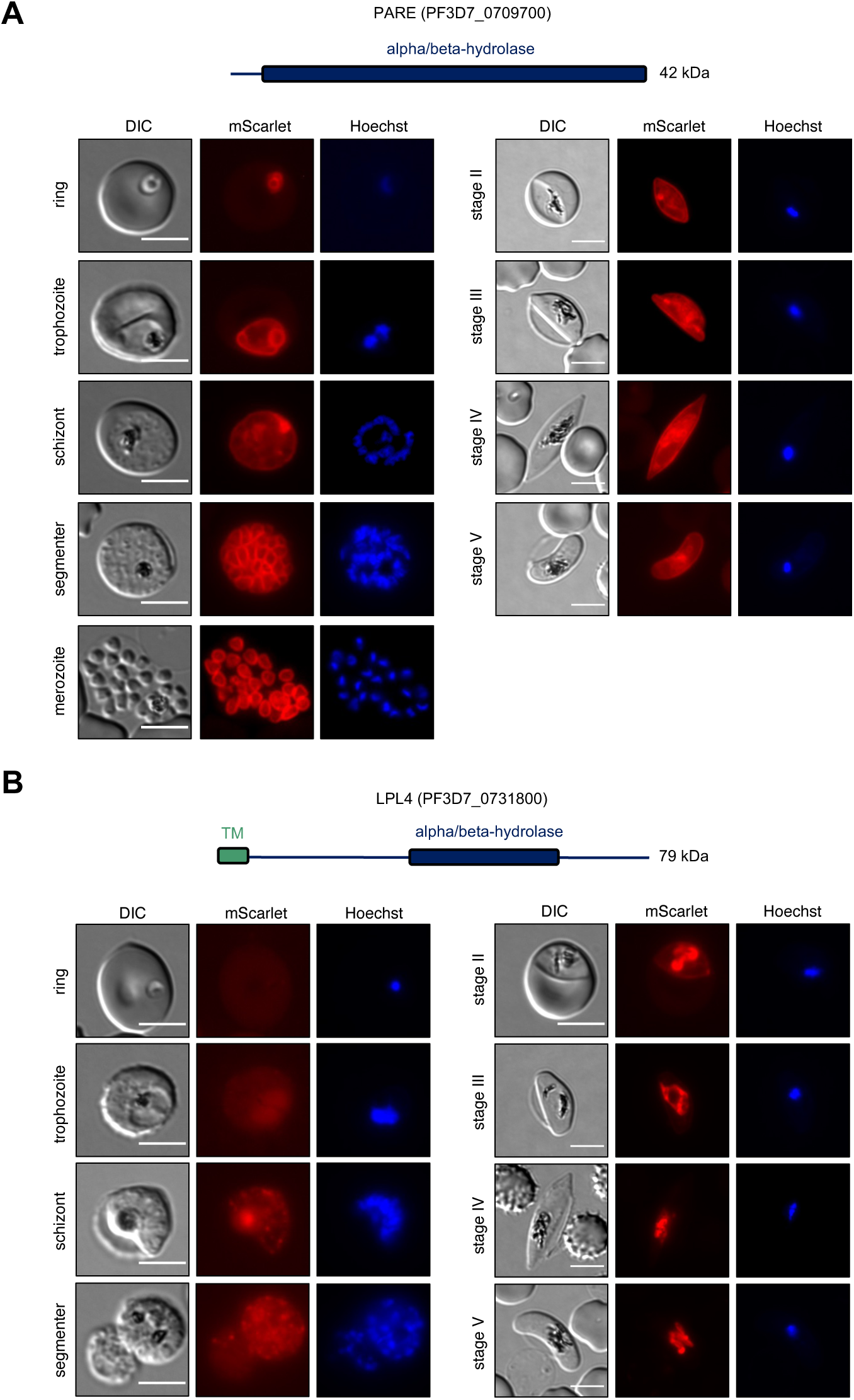
Localization analysis of PARE and LPL4. Parasites expressing endogenously tagged PARE-mScarlet (A) and LPL4-mScarlet (B) were analyzed during asexual and sexual blood stage development by live-cell microscopy. Nuclei were stained with Hoechst. Scale bars = 5 µm. DIC, differential interference contrast. Schematic representations of the functional domains of the two proteins are shown on top of the images. TM, transmembrane domain.

**FIGURE 3.**
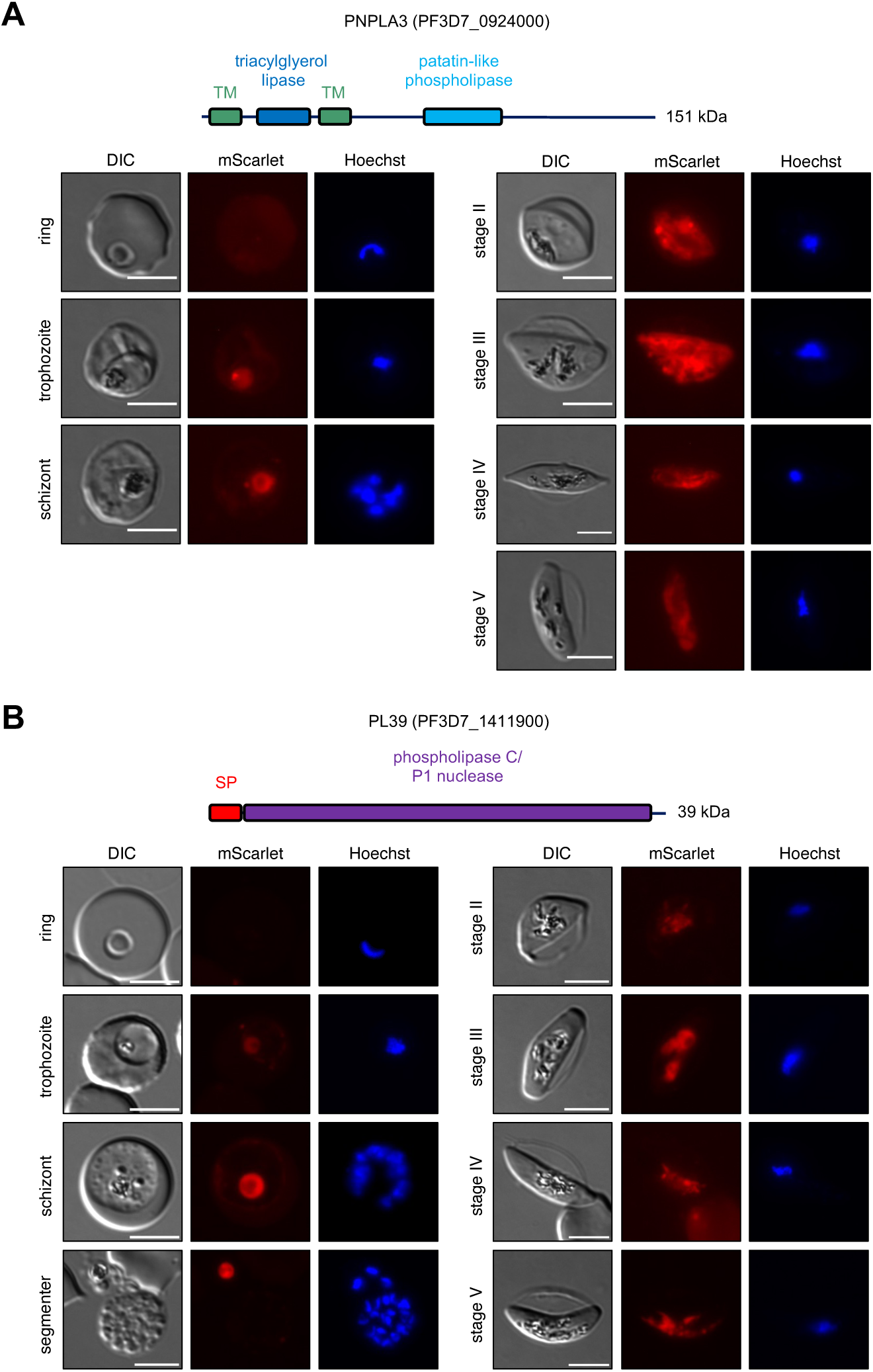
Localization analysis of PNPLA3 and PL39. Parasites expressing endogenously tagged PNPLA3-mScarlet (A) and PL39-mScarlet (B) were analyzed during asexual and sexual blood stage development by live-cell microscopy. Nuclei were stained with Hoechst. Scale bars = 5 µm. DIC, differential interference contrast. Schematic representations of the functional domains of the two proteins are shown on top of the images. SP, signal peptide; TM, transmembrane domain.

**FIGURE 4.**
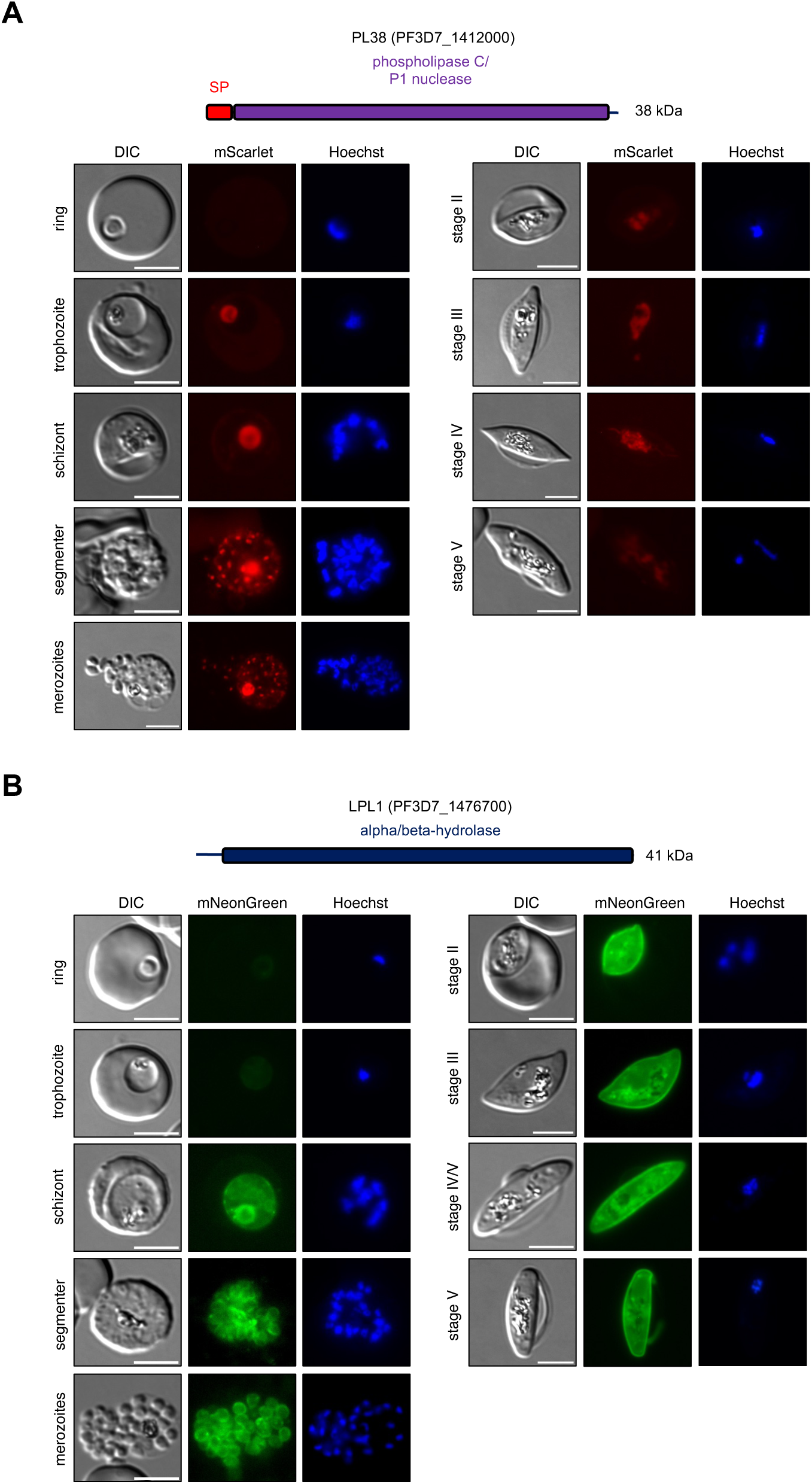
Localization analysis of PL38 and LPL1. Parasites expressing endogenously tagged PL38-mScarlet (A) and LPL1-mNeonGreen (B) were analyzed during asexual and sexual blood stage development by live-cell microscopy. Nuclei were stained with Hoechst. Scale bars = 5 µm. DIC, differential interference contrast. Schematic representations of the functional domains of the two proteins are shown on top of the images. SP, signal peptide.

The lysophospholipase 1 (LPL1, PF3D7_1476700) (Asad *et al*., 2021) fused to mScarlet showed a peripheral localization in gametocytes, while no signal was visible in asexual blood stages of this parasite line (Figure S3). Given that evidence for expression of LPL1 in asexual blood stages was provided previously (Asad *et al*., 2021), we generated an additional parasite line, in which the gene was C-terminally tagged with the in comparison to mScarlet much brighter green fluorescent protein mNeonGreen (Shaner *et al*., 2013). This genetic modification was performed in a 3D7 cell line that fully supports gametocyte development (Filarsky *et al*., 2018) and successful integration of the targeting construct was confirmed by PCR (Figure S1). LPL1-mNeonGreen displayed a strong peripheral signal throughout gametocyte development. In contrast to LPL1-mScarlet, a weaker peripheral signal was also observed for LPL1-mNeonGreen in schizonts as well as free merozoites (Figure 4B). This suggests that LPL1 localizes to the PPM and that its apparently lower expression levels in asexual blood stages prevented its detection in the LPL1-mScarlet parasite line.

### Loss of individual phospholipases does not impair gametocyte development

To probe into the physiological function of the six putative phospholipases for gametocyte development and egress, we next performed targeted gene disruption using the SLI system (Birnbaum *et al*., 2017) in the NF54/iGP2 background (Boltryk *et al*., 2021). Correct integration of the respective targeting constructs was confirmed by PCR (Figure S4). Previously, five of the six phospholipases (PARE, LPL4, PNPLA3, PL39, and PL38) were demonstrated to be non-essential for asexual blood stage development using the same targeting constructs (Burda *et al*., 2021). We thus only analyzed asexual blood stage proliferation of LPL1-knockout (KO) parasites. This, however, did not reveal an impaired growth in the presence or absence of 2 mM choline as compared to wildtype (WT) parasites, suggesting that LPL1 plays a non-essential or redundant function in asexual blood stage parasites (Figure S5).

To investigate the function of the selected phospholipases for gametocyte development, we induced gametocyte commitment as previously described (Boltryk *et al*., 2021) and followed gametocyte development over two weeks. Gametocyte survival rates, calculated by the ratio of gametocytemia on day 11 and day 5 of gametocyte development, were not significantly different between WT and KO-parasite lines (Figure S6). In line with this, quantification of gametocyte stages over the 11-day course of gametocytogenesis of all phospholipase-KO lines revealed no major defect in gametocyte maturation (Figure 5). Taken together, these data suggest that the six analyzed putative phospholipases are dispensable for gametocyte development.

**FIGURE 5.**
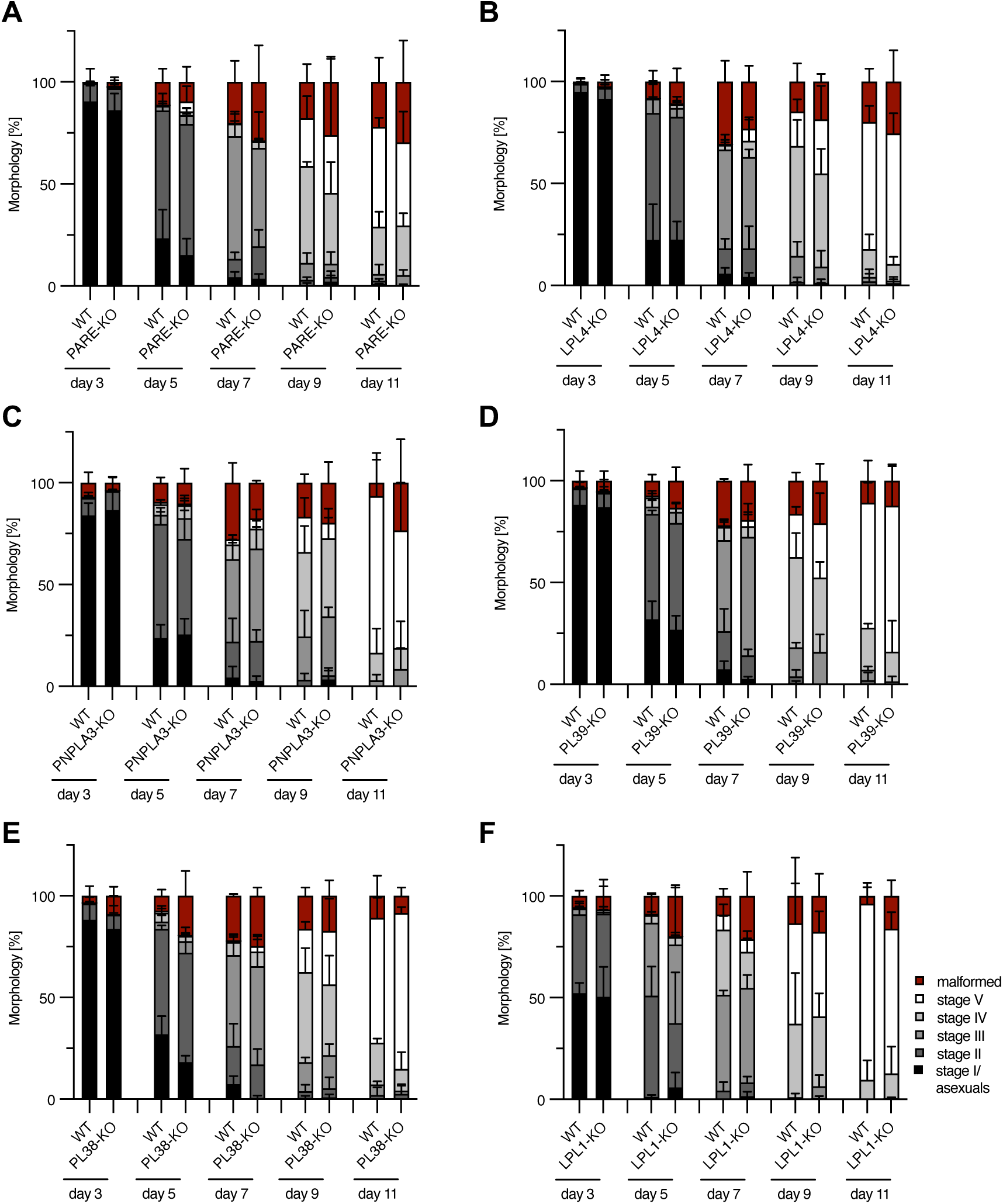
Disruption of the six individual genes encoding for putative phospholipases does not affect gametocyte maturation. A-F) Quantification of gametocyte stages on the days of gametocyte development indicated below the graphs by counting Giemsa-stained thin blood smears. Shown are means + SD of three independent experiments. In some experiments, NF54/iGP2 WT gametocytes served as a control for several KO lines in parallel. (A) PARE-KO, (B) LPL4-KO, (C) PNPLA3-KO, (D) PL39-KO, (E) PL38-KO and (F) LPL1-KO.

### Disruption of PNPLA3 delays male gametocyte exflagellation

Next, we analyzed gamete egress from mature stage V gametocytes. During this process, male gametocytes undergo a striking transformation resulting in the formation of eight haploid motile microgametes, while female gametes maintain their spherical shape after egress from the RBC (Kuehn and Pradel, 2010). To study whether gametocyte egress is affected in the KO parasite lines, we used an established egress assay based on live-cell fluorescence microscopy (Figure 6A,B) (Suareź-Cortés *et al*., 2014). For this, the RBC membrane of RBCs infected with mature stage V gametocytes was stained using the live-cell dye iFluor555-wheat germ agglutinin (WGA) before activation. Subsequently, gametocytes were activated for 20 minutes and parasite morphology (falciform shape versus spherical shape) as well as the WGA staining pattern (WGA-positive versus WGA-negative) were investigated (Suareź-Cortés *et al*., 2014). As expected, the vast majority of non-activated gametocytes showed a falciform shape with strong WGA staining of the erythrocyte membrane, while after activation most gametocytes displayed a round morphology and were WGA-negative, indicating successful egress from the host RBC (Figure 6C-H). Hereby, no major differences between WT and KO parasite lines were observed, indicating that all of the six tested phospholipases are dispensable for gamete egress. To further analyze egress, we also separately quantified exflagellation rates of male gametocytes on four subsequent days from day 11 until day 14 of gametocyte development. To that end, we activated gametocytes for 12 minutes and subsequently determined the percentage of exflagellating gametocytes within the next 5 minutes. PARE-KO and LPL1-KO gametocytes showed similar exflagellation rates to WT parasites, while LPL4-KO, PL39-KO and PL38-KO showed significantly higher exflagellation rates compared to WT parasites (Figure 7). In line with the WGA staining-based egress assay, we can thus conclude that none of these five putative phospholipases plays an essential role in gametocyte exflagellation. Remarkably, PNPLA3-KO gametocytes showed a significant reduction in exflagellation within the initial five minutes of observation time. Extending observation time by 10 minutes resulted in exflagellation rates similar to WT parasites (Figure 7C), suggesting that loss of PNPLA3 delays efficient exflagellation but does not prevent this process from happening.

**FIGURE 6.**
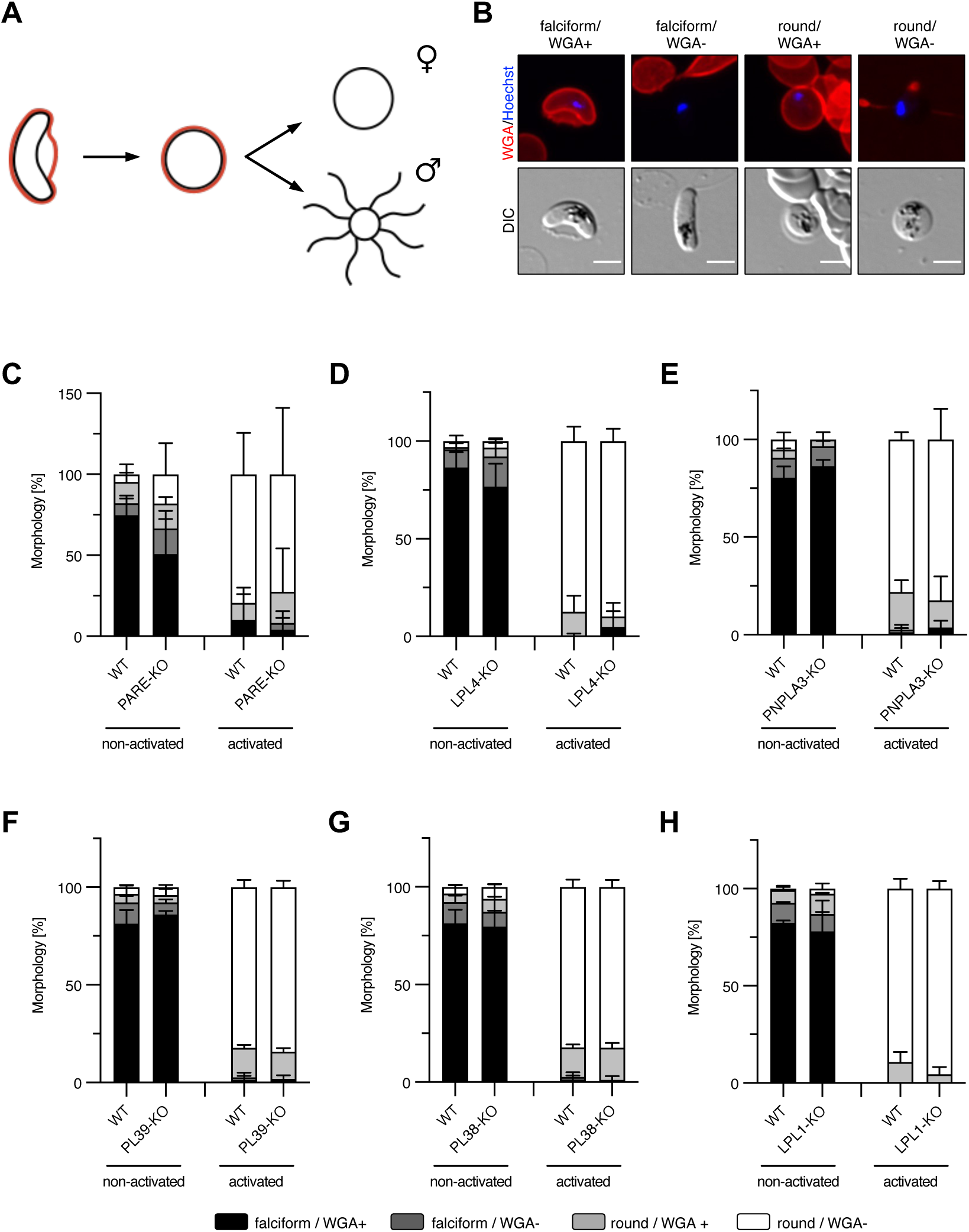
None of the putative phospholipase-KO lines displays a defect in gamete egress. A) Schematic of the gamete egress assay. On day 14 of gametocytogenesis, the RBC membrane was stained with iFluor555-WGA (red). Upon activation, gametocytes first round up before egressing from the RBC thereby losing the iFluor555-WGA signal. Female gametocytes retain a round shape, while male gametocytes form eight motile flagella. B) Representative images of the four categories used to classify gametocytes/gametes in the gamete egress assay. Staining with iFluor555-WGA (red) and Hoechst (blue). DIC, differential interference contrast. Scale bars = 5 µm. C-H) Imaging-based gamete egress assays with (C) PARE-KO, (D) LPL4-KO, (E) PNPLA3-KO, (F) PL39-KO, (G) PL38-KO, and (H) LPL1-KO gametocytes. Gametocytes were imaged non-activated and after activation for 20 min. At least 20 parasites were imaged per condition. Shown are means + SD of three independent experiments.

**FIGURE 7.**
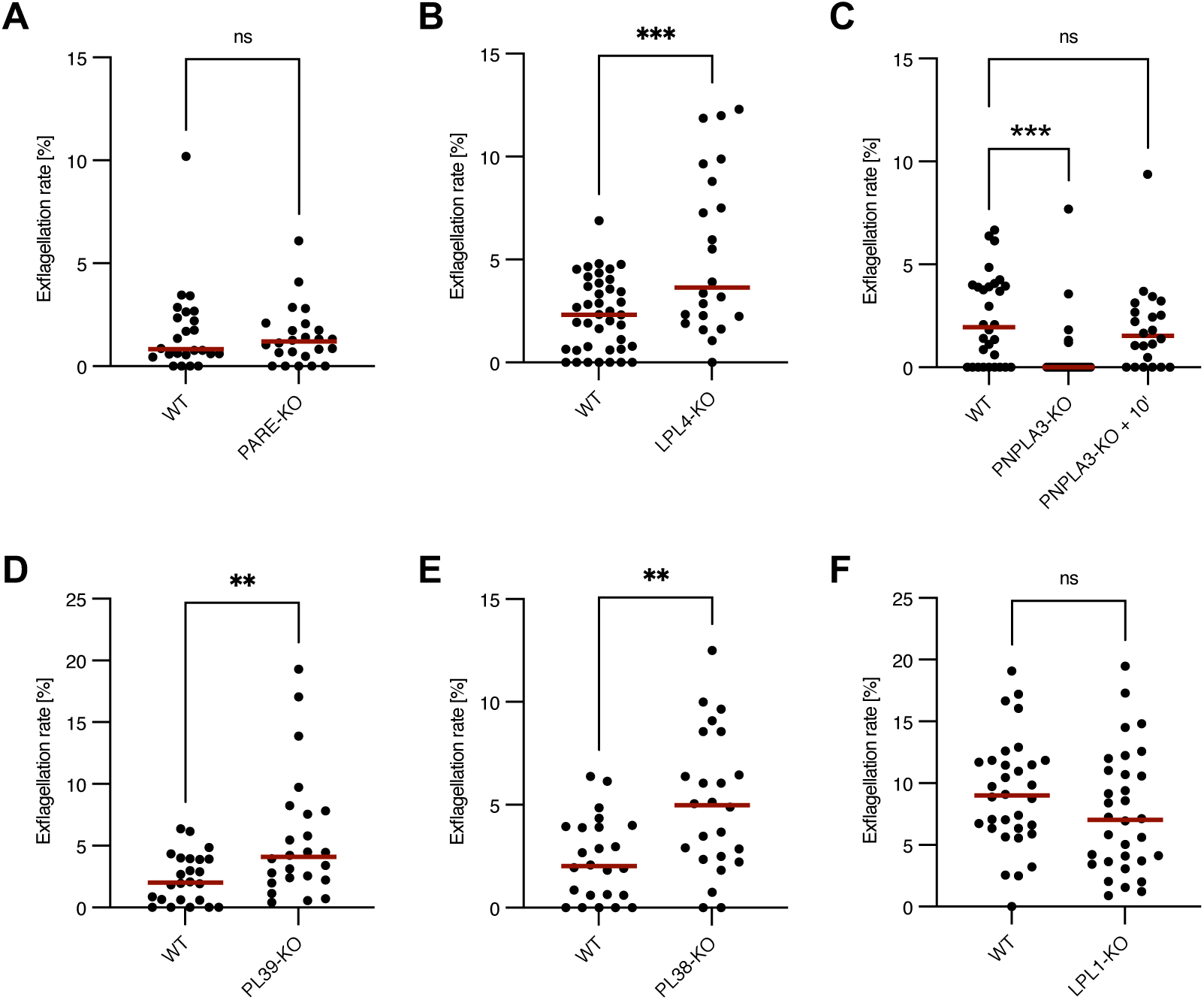
PNPLA3-KO parasites show a delay in male gametocyte exflagellation. A-F) Exflagellation assays of (A) PARE-KO, (B) LPL4-KO, (C) PNPLA3-KO (includes data with 10 additional minutes of exflagellation time), (D) PL39-KO, (E) PL38-KO, and (F) LPL1-KO gametocytes. Graphs display superplots of at least three independent experiments. For each experiment, exflagellation was checked in technical duplicates on four subsequent days (day 11 – day 14 of gametocyte development). Red lines indicate the median of all exflagellation rates observed across all days and experiments for the respective parasite line. For statistical analysis of the results, an unpaired students t-test was used (**p < 0.01; ***p < 0.0005; n.s., not significant). In some experiments, NF54/iGP2 gametocytes served as control for several KO lines in parallel.

## DISCUSSION

Phospholipases were shown to have important functions for asexual intraerythrocytic development and egress of malaria blood and liver stage parasites (Burda *et al*., 2015; Burda *et al*., 2021; Ramaprasad *et al*., 2023). We here aimed to expand the functional characterization of these enzymes and investigated the role of six putative phospholipases for gametocytogenesis and gametocyte egress.

Two putative phospholipases analyzed in this study were PARE and LPL1. PARE was previously reported to have esterase activity and to activate esterified pepstatin, a peptidyl inhibitor of malarial aspartyl proteases, although its localization in the parasite had not yet been determined (Istvan *et al*., 2017). By endogenous tagging, we now localized PARE and LPL1 to the PPM in asexual and sexual blood stages of the parasite. In addition to this, we showed that disruption of PARE and LPL1 does not affect asexual blood stage development (this study and (Burda *et al*., 2021)), arguing for a redundant function of both enzymes under *in vitro* conditions. These localization and functional data regarding LPL1 differ to a previous study that localized LPL1 to a multi-vesicular neutral lipid-rich body next to the food vacuole in asexual blood stages and that proposed an essential role of LPL1 in neutral lipid synthesis contributing to hemozoin formation in the parasite (Asad *et al*., 2021). While the reasons for the observed differences in LPL1 localization remain unclear, the absence of a growth defect in asexual blood stages due to LPL1 disruption could be related to the different methods used for the inactivation of LPL1. While Asad *et al*. employed a conditional degradation approach for ablating LPL1 expression, we used targeted gene disruption in this study involving selection of transgenic parasites over several intraerythrocytic cycles that may possibly enable parasites to adapt to the loss of LPL1 through a currently unknown mechanism.

Two other putative phospholipases that we analyzed in this study were PL39 and PL38, which both contain predicted signal peptides and phospholipase C/P1 nuclease domains. Enzymes containing these domains are typically involved in phosphate ester hydrolysis of lipids or nucleic acids (Coleman, 1992; Desai and Shankar, 2003). The genomic proximity of *pl39* and *pl38* and the existence of only one single orthologue in the rodent malaria parasite *P. berghei* at this genomic location (according to PlasmoDB.org) indicates that these loci might have arisen from gene duplication. However, as PL39 and PL38 display only 46 % sequence identity (according to ClustalW), they may have diverged to serve distinct functions over the course of evolution. In line with this, PL39 localized to the food vacuole, while PL38 displayed a possible apical localization. Interestingly, the *Toxoplasma gondii* patatin-like phospholipase *Tg*PL3 (TGME49_305140) has been localized to the apical pole where its phospholipase activity is necessary for rhoptry secretion (Wilson *et al*., 2020). We speculate that the putative phospholipase activity of PL38 might similarly contribute to invasion. However, gene deletion did not affect asexual blood stage proliferation, suggesting an unimpaired invasion process of PL38-deficient parasites (Burda *et al*., 2021).

To explore the role of the six selected putative phospholipase candidates for gametocytogenesis and gametocyte egress, we performed targeted gene disruption of the individual genes. We then analyzed sexual blood stage development of the resulting mutant parasite lines. This revealed that gametocyte survival and maturation were not affected by the individual disruption of the six selected putative phospholipases, suggesting that, similar to our results obtained in asexual blood stages (Burda *et al*., 2021), a high level of functional redundancy also exists among the parasite phospholipases during sexual blood stage development.

A live-cell fluorescence microscopy-based egress assay using the RBC membrane stain WGA (Suareź-Cortés *et al*., 2014) did not reveal any substantial differences in gametocyte egress between WT and mutant parasite lines. Interestingly, however, separately quantifying exflagellation rates of male gametocytes revealed that parasites lacking the PNPLA3 showed a significant delay in efficient exflagellation of male gametocytes indicating a critical, yet not essential, role of this enzyme in male gametogenesis. It is important to note that exflagellation rates of PNPLA3-null parasites reached WT levels, when the analysis time was extended for 10 minutes. Since activation of gametocytes for the WGA assay was done for 20 minutes, while 12 minutes of gametocyte activation were used for the exflagellation assay, the extended time for gametocyte activation could explain why the PNPLA3-associated phenotype in exflagellation was not observed in the WGA-based egress assay. What specific role PNPLA3 might play in the exflagellation process remains to be elucidated. Endogenous tagging of PNPLA3 revealed a vesicular and diffuse cytoplasmic localization pattern arguing against a direct function of PNPLA3 in disruption of the PVM or the host cell membrane. Notably, conditional inactivation of the cytoplasmic PNPLA1 (PF3D7_0209100) reduced efficiency of gametocyte egress possibly by interfering with the release of egress-associated vesicles (Singh *et al*., 2019). It is thus reasonable to hypothesize that a similar function might also be performed by PNPLA3. Given that conditional inactivation of PNPLA1 only resulted in a partial egress phenotype, both enzymes might even work together during egress, a scenario that should be explored in future studies by generating mutants that lack both enzymes.

Taken together, our data suggest that all six phospholipase candidates may have redundant functions for gametocyte development and gamete egress. However, PNPLA3 might have a role in efficient exflagellation of male gametocytes, identifying it as a novel player in this important process for transmission of the parasite.

## MATERIALS AND METHODS

### Cloning of SLI-based targeting constructs

For the generation of SLI-based tagging constructs, we exchanged the GFP coding sequence of pSLI-GFP (Birnbaum *et al*., 2017) by mScarlet. For this, we first generated the intermediate construct pSLI-PF3D7_0924000-GFP by amplifying the C-terminal 973 bp of the coding sequence by PCR using primers PF3D7_0924000-TAG-fw/ PF3D7_0924000-TAG-rev, starting with a stop codon, to serve as homology regions for single-crossover based integration, followed by cloning the PCR product using NotI/MluI into pSLI-GFP. Subsequently, we amplified the mScarlet coding sequence from SP-mScarlet (Mesén-Ramírez *et al*., 2019) using primers mScarlet-fw/mScarlet-rev and cloned it via AvrII/SalI into pSLI-PF3D7_0924000-GFP, thereby replacing the GFP cassette. This resulted in the final targeting plasmid pSLI-PF3D7_0924000-mScarlet. For generation of the other mScarlet-tagging constructs, the respective C-terminal targeting sequences were amplified by PCR, starting with a stop codon, and cloned using NotI/MluI into pSLI-PF3D7_0924000-mScarlet.

For generation of the pSLI-PF3D7_1476700-mNeonGreen-glmS construct, the 963 bp C-terminal region of *pf3d7_1476700* was amplified by PCR using primers PF3D7_1476700-TAG-fw/ PF3D7_1476700-TAG-rev and cloned into pSLI-mNeonGreen-glmS (kindly provided by Arne Alder) using NotI/MluI. The pSLI-mNeonGreen-glmS vector is based on pSLI-GFP-GlmS (Burda *et al*., 2020) but contains the mNeonGreen coding sequence (Shaner *et al*., 2013) instead of the one of GFP.

For the generation of the SLI-based KO construct pSLI-PF3D7_1476700-TGD, 423 bp immediately downstream of the start ATG of the *lpl1* gene were amplified by PCR using primers PF3D7_1476700-KO-fw/ PF3D7_1476700-KO-rev and cloned using NotI/MluI into pSLI-GFP (Birnbaum *et al*., 2017) to generate the final targeting plasmid. The generation of all other employed pSLI-based KO constructs was described previously (Burda *et al*., 2021).

Phusion High-Fidelity DNA polymerase (New England BioLabs) was used for all plasmid constructions and all plasmid sequences were confirmed by Sanger sequencing. For sequences of oligonucleotides used in this study see Supplemental file 1.

### P. falciparum culture

Blood stages of 3D7 and NF54 *P. falciparum* parasites and transgenic derivates were cultured in human RBCs. Cultures were maintained at 37°C in an atmosphere of 94% nitrogen, 5% carbon dioxide and 1% oxygen using RPMI complete medium containing 0.5% Albumax according to standard procedures (Trager and Jensen, 1976). If not stated otherwise, medium was supplemented with 2 mM choline. Cultures of NF54/iGP2-based parasites were cultured in the presence of 2.5 mM glucosamine (GlcN) to prevent overexpression of the gametocyte commitment factor GDV1 (Boltryk *et al*., 2021).

### Generation of SLI-based parasite lines

For transfection of constructs, Percoll (GE Healthcare)-enriched synchronized mature schizonts were electroporated with 50 μg of plasmid DNA using a Lonza Nucleofector II device (Moon *et al*., 2013). Transfectants were selected in medium supplemented with 3 nM WR99210 (Jacobus Pharmaceuticals). For generation of stable integrant cell lines, parasites containing the episomal plasmids selected with WR99210 were grown with 400 μg/ml Neomycin/G418 (Sigma) to select for transgenic parasites carrying the desired genomic modification as described previously (Birnbaum *et al*., 2017). Successful integration was confirmed by diagnostic PCR using FIREpol DNA polymerase (Solis BioDyne). For primer sequences see Supplemental file 1.

### Production of synchronous *P. falciparum* gametocytes

Gametocytes of NF54/iGP2-derived parasites were basically induced as previously described (Boltryk *et al*., 2021). For this, synchronous ring stage cultures (2-3 % parasitemia) were washed in medium to remove remaining GlcN and plated at 2.5-5% hematocrit in culture medium without GlcN (= day –1 of gametocyte development). After reinvasion of committed rings (= day 1 of gametocyte development), asexual parasites were depleted using 50 mM N-acetyl-D-glucosamine (GlcNAc) in the culture medium. GlcNAc was added to the cell culture medium until day 6. From day 7 until day 14, gametocyte cultures were fed with culture medium containing 0.25 % Albumax + 5 % human serum. Gametocytes were cultured in the presence of GlcN from day 1 onwards and gametocyte cultures were fed daily.

Induction of 3D7-derived gametocytes was performed as previously described (Filarsky *et al*., 2018). To this aim, synchronous ring stage cultures were washed twice and then cultivated in medium without choline (= day –1 of gametocyte development). From day 1 of gametocyte development until day 6, asexual parasites were depleted using 50 mM GlcNAc in the culture medium (now again containing choline). From day 7 until day 14, gametocyte cultures were fed with culture medium containing 0.25 % Albumax + 5 % human serum and choline. Gametocyte cultures were fed daily. Gametocyte stages and gametocytemia were monitored using Giemsa-stained thin blood smears.

### Fluorescence microscopy

For staining of nuclei, parasites were incubated with 0.45 µg/ml Hoechst in culture medium for 30 min at 37°C. Images were acquired on a Leica D6B fluorescence microscope, equipped with a Leica DFC9000 GT camera and a Leica Plan Apochromat 100×/1.4 oil objective. Image processing was performed using ImageJ and representative images were adjusted for brightness and contrast.

### Growth analysis of mutant parasites

For asexual blood stage growth analysis of parasites, schizont stage parasites were isolated by Percoll enrichment and incubated with uninfected RBCs (5% hematocrit) for 3 hours to allow rupture and invasion. Parasites were then treated with 5% sorbitol to remove residual unruptured schizonts, leading to a synchronous ring stage culture with a 3-hour window. These were allowed to mature to trophozoites for one day and parasitemia was determined by flow cytometry and adjusted to exactly 0.1% starting parasitemia in a 2 ml dish. Medium was changed daily and growth of the parasite lines was assessed by flow cytometry over three intraerythrocytic cycles when parasites were in the trophozoite stage. As a reference, WT parasites were included in each assay.

### Flow cytometry

Flow cytometry-based analysis of parasite lines was performed essentially as described previously (Malleret *et al*., 2011). In brief, 20 μl resuspended parasite culture were incubated with dihydroethidium (5 μg/ml, Cayman) and SYBR Green I dye (0.25 x dilution, Invitrogen) in a final volume of 100 μl medium for 20 min at room temperature protected from light. Samples were analyzed on a ACEA NovoCyte flow cytometer. RBCs were gated based on their forward and side scatter parameters. For every sample, 100,000 events were recorded and parasitemia was determined based on SYBR Green I fluorescence.

### Imaging-based gametocyte egress assay

On day 14 of gametocyte development, the egress of mature male and female gametocytes was tested as described previously (Suareź-Cortés et al., 2014) with minor modifications. In brief, parasite-infected RBC were stained with iFluor555-WGA (Biomol, stock 2 mg/ml, final concentration 5 µg/ml) and 0.45 µg/ml Hoechst at 37°C. After 30 min, the samples were washed in prewarmed Ringer solution. 5 µL were placed on a slide and immediately covered with a cover slip. Imaging of this non-activated control was performed for 20 min. To study egress, gametocytes were activated by incubation in ookinete medium for 20 min at 26 °C before imaging for 20 min at room temperature. Ookinete medium was prepared by supplementing culture medium (without serum or albumax) with 100 µM Xanthurenic acid and adjusting the pH to 8.0, followed by addition of 20 % human serum.

### Gametocyte exflagellation assay

From day 11 until day 14 of gametocytogenesis, exflagellation was determined daily in technical duplicates. 250 µl of resuspended gametocyte culture were spun down in an Eppendorf tube at 800 x g for 1 min. The supernatant was discarded and the pellet was resuspended in 250 µl prewarmed (26 °C) ookinete medium. Gametocytes were incubated for 12 min at 26 °C. Then, 6 µl were placed on a slide and covered with a coverslip. For 5 min, all gametocytes as well as exflagellation events were counted using a 40× oil objective (aperture closed as much as possible to contrast the hemozoin). If indicated, the observation time was extended for another 10 min to detect a potential delay in exflagellation in non-exflagellating mutants.

### Statistical analysis

For statistical analysis of differences between two groups, unpaired two-tailed students t-tests were used. All statistical tests were done in GraphPad Prism. P values of <0.05 were considered significant. Statistical details (n numbers, test used, definition of the error bars) are described in the figure legends.

## DATA AVAILABILITY

All data generated or analyzed during this study are included in this published article and its supplemental material files.

## ACKNOWLEDGEMENTS

We thank Danny Wilson for critical reading of the manuscript and Arne Alder for providing the pSLI-mNeonGreen-glmS vector. We acknowledge the Advanced Light and Fluorescence Microscopy (ALFM) facility at the Centre for Structural Systems Biology (CSSB) for support with image recording and analysis. This work was supported by a grant from the German Research Foundation (Deutsche Forschungsgemeinschaft, DFG) to PCB (project number 414222880). KN was funded by a grant from the DFG within the SPP2225 to TWG and PCB (project number 446556156).

## AUTHOR CONTRIBUTIONS

Conceived and designed the experiments: EP, PCB. Performed the experiments: EP, KN, PCB. Analyzed the data: EP, PCB, KN, TWG. Paper writing: PCB, EP, KN, TWG and based on the PhD thesis of EP.

## SUPPLEMENTARY INFORMATION

**FIGURE S1.**
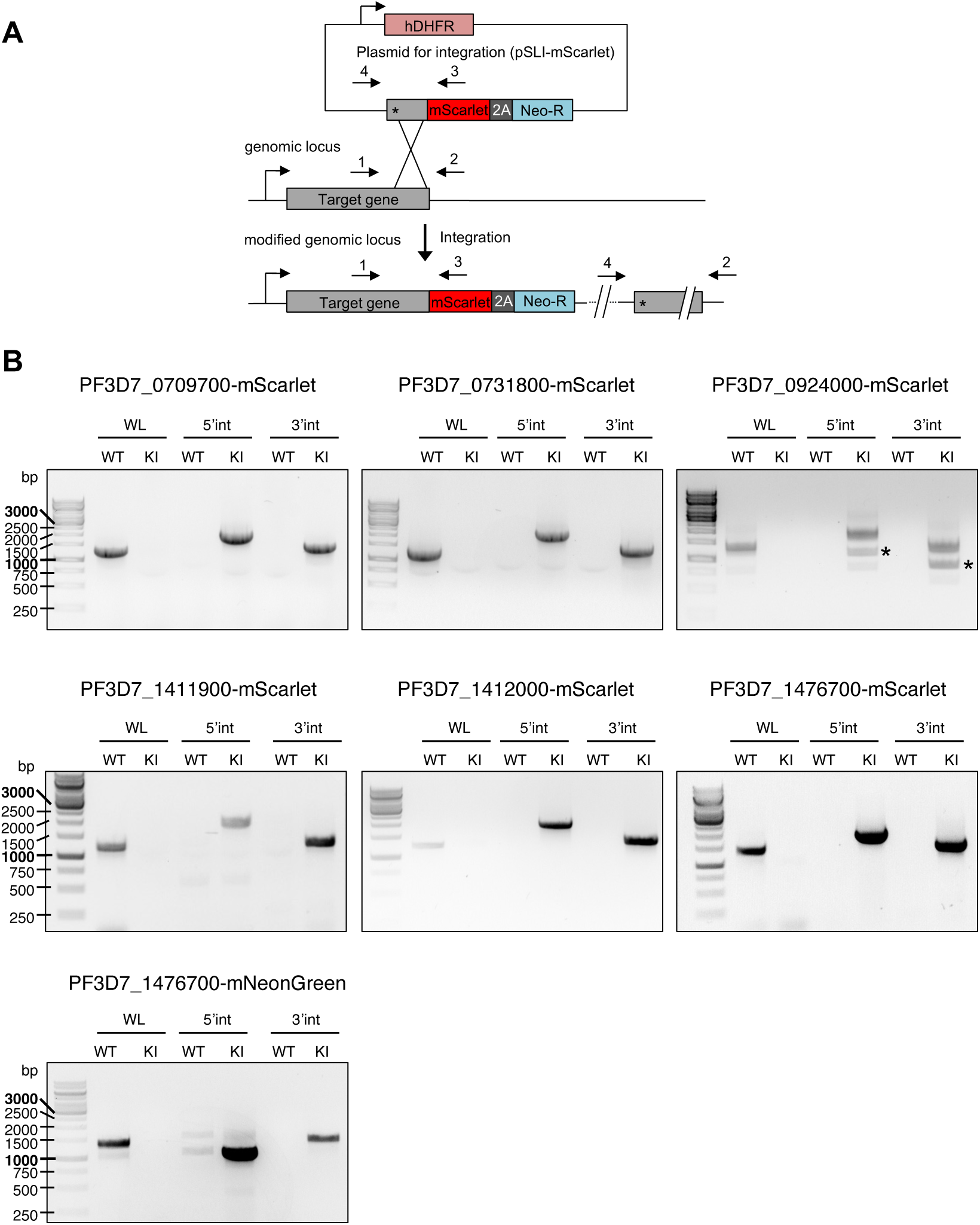
Analytical PCRs of mScarlet/mNeonGreen-tagged parasite lines in the NF54/iGP2 and 3D7 background. A) Schematic representation of the selection-linked integration (SLI) strategy used for endogenous mScarlet tagging is shown. A similar strategy was used for endogenous mNeonGreen tagging. Binding sites of primers used to detect successful integration of the targeting constructs by PCR are indicated. 2A, skip peptide; Neo-R, neomycin-resistance gene; asterisks, stop codons. B) Agarose gel electrophoresis of PCR products amplified from genomic DNA of transgenic mScarlet/mNeonGreen-tagged parasite lines as well as unmodified wildtype (WT) parasites. WL, WT locus (primer 1+2); 5’int, 5’ integration (primer 1+3); 3’int, 3’ integration (primer 4+2); KI, knock in cell line; *, unspecific double-band.

**FIGURE S2.**
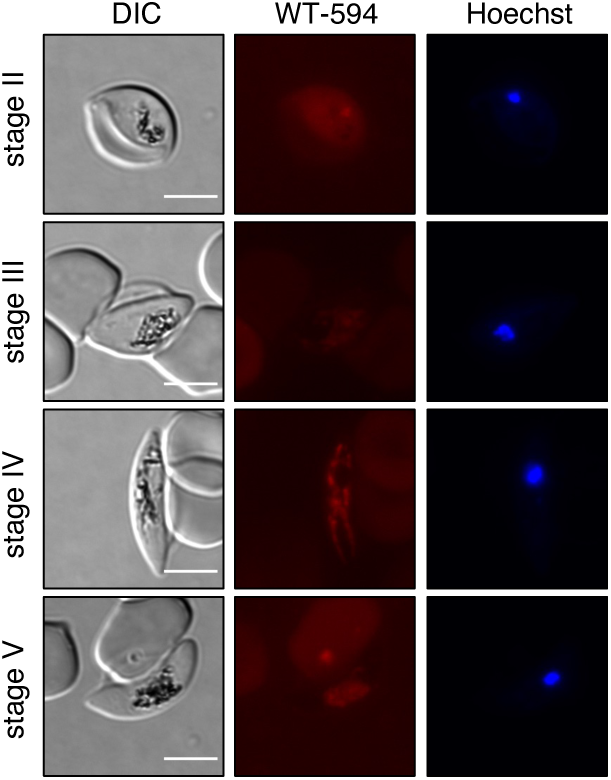
Background fluorescence in live-cell microscopy of NF54/iGP2 WT gametocytes. NF54/iGP2 WT gametocytes were imaged with the same imaging settings used for analysis of transgenic parasite lines expressing mScarlet fusion proteins. Nuclei were stained with Hoechst. Scale bars = 5 µm. DIC, differential interference contrast.

**FIGURE S3.**
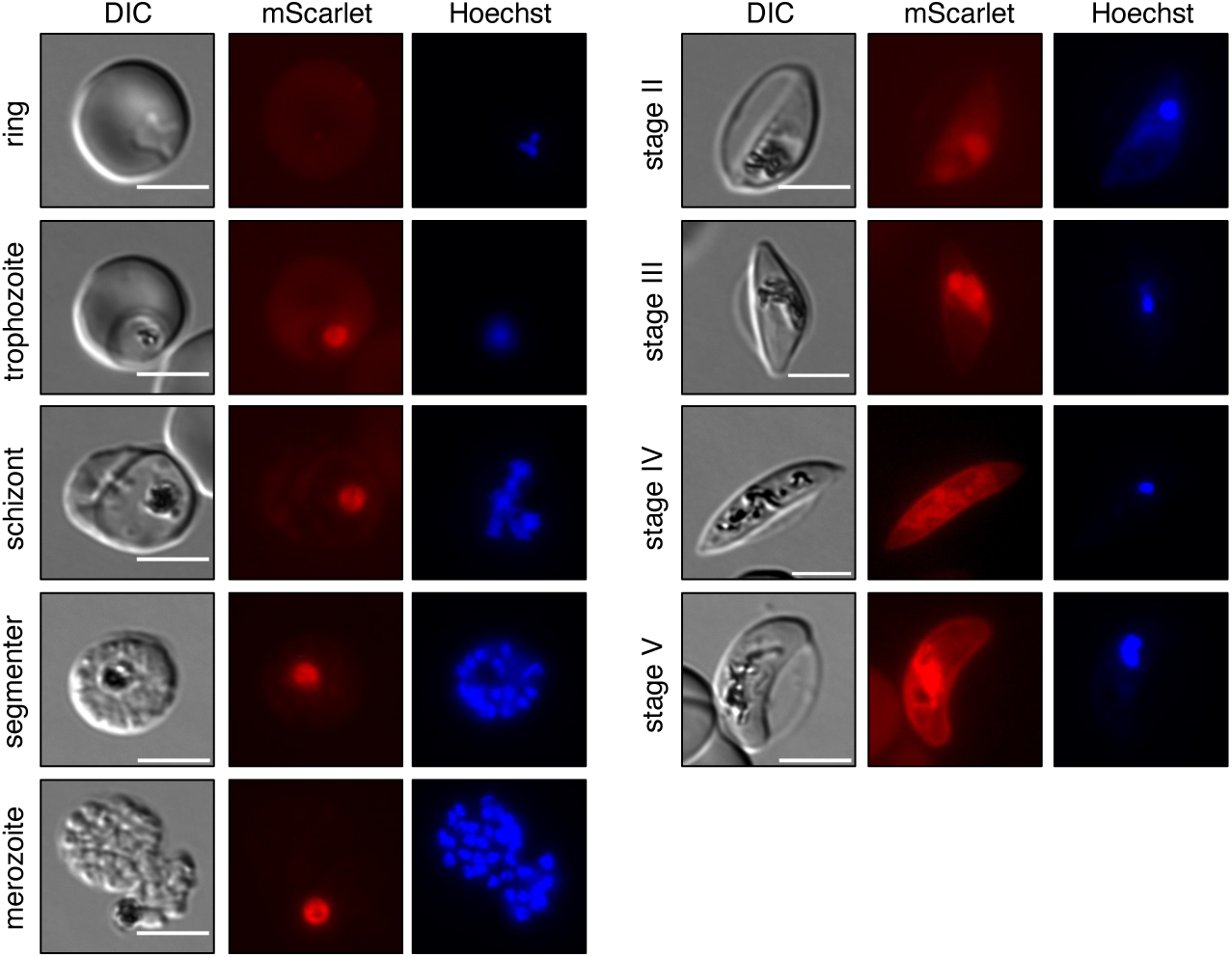
Live-cell microscopy of LPL1-mScarlet parasites during asexual and sexual blood stage development. Nuclei were stained with Hoechst. Scale bars = 5 µm. DIC, differential interference contrast.

**FIGURE S4.**
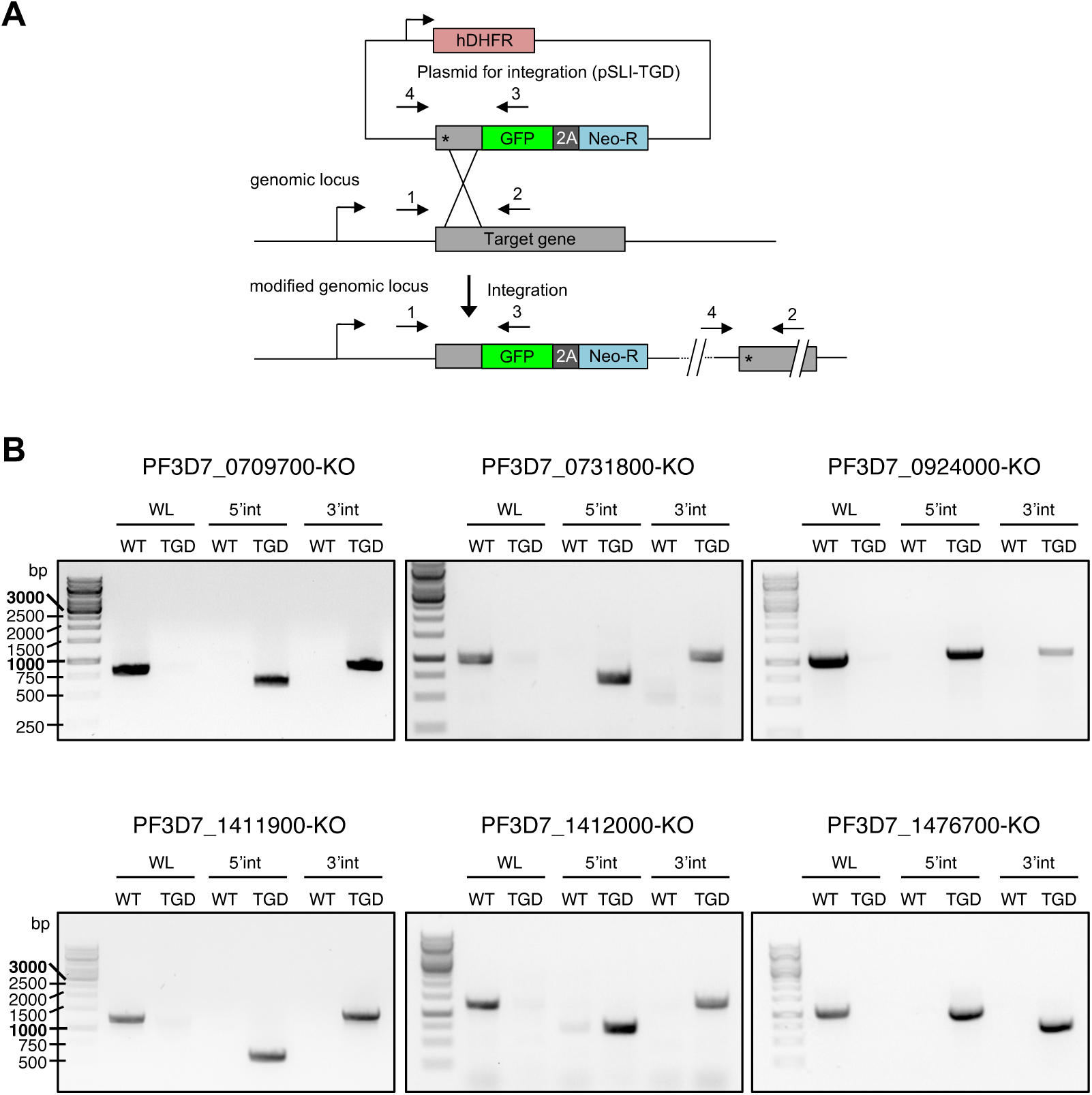
Analytical PCRs of KO parasite lines in the NF54/iGP2 background. A) Schematic representation of the selection-linked integration (SLI) strategy used for targeted gene disruption is shown. Binding sites of primers used to detect successful integration of the targeting constructs by PCR are indicated. 2A, skip peptide; Neo-R, neomycin-resistance gene; asterisks, stop codons. B) Agarose gel electrophoresis of PCR products amplified from genomic DNA of transgenic KO parasite lines as well as unmodified wildtype (WT) parasites. WL, WT locus (primer 1+2); 5’int, 5’ integration (primer 1+3); 3’int, 3’ integration (primer 4+2).

**FIGURE S5.**
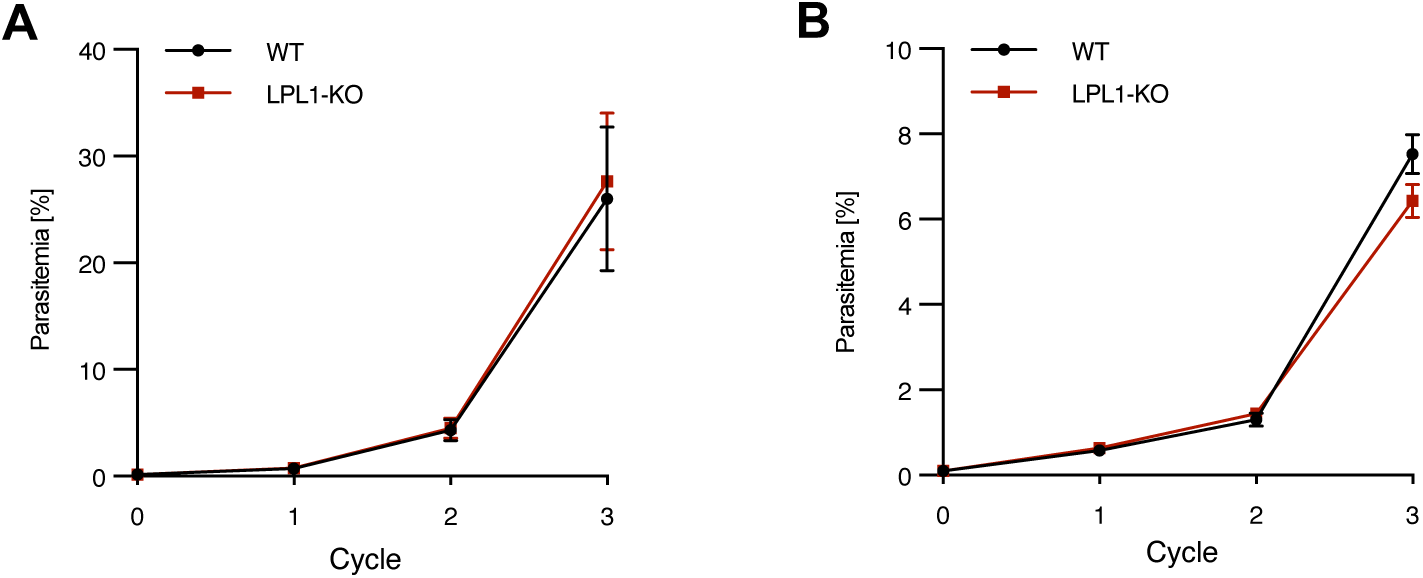
Growth assay of LPL1-KO parasites over three intraerythrocytic cycles in the presence (A) or absence (B) of 2 mM choline. Parasites were diluted 1:10 after the second cycle to prevent overgrowth. Shown are means +/-SD of three independent experiments. Unpaired students t-test revealed no significant differences between WT and KO parasites.

**FIGURE S6.**
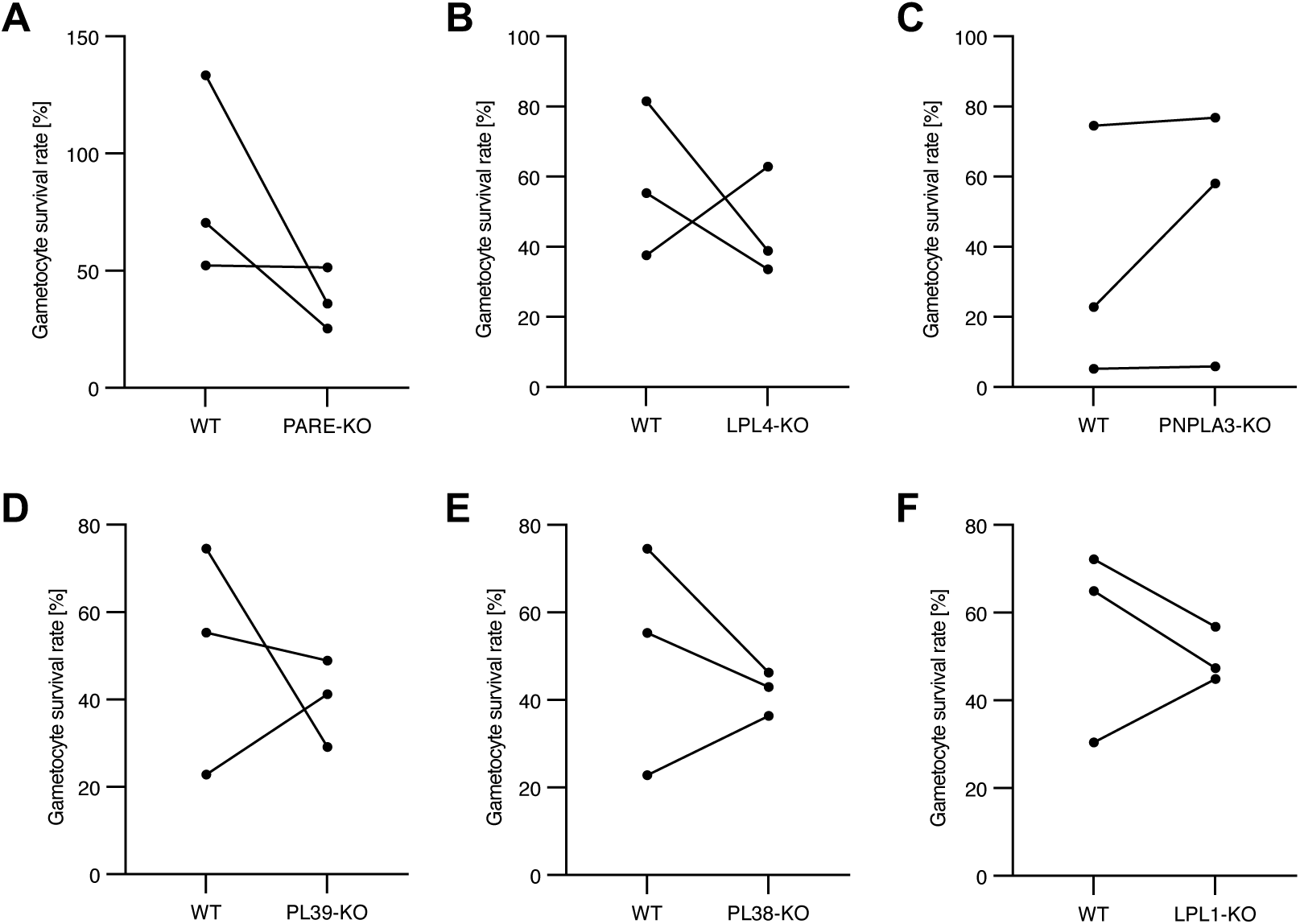
Disruption of the six individual genes encoding for putative phospholipases does not affect gametocyte survival. (A-F) Gametocyte survival rates as calculated by the ratio of gametocytemia on day 11 and day 5 of gametocyte development. Dots represent independent experiments. Lines connect the KO with the respective WT control. In some experiments, NF54/iGP2 WT gametocytes served as a control for several KO lines in parallel. (A) PARE-KO, (B) LPL4-KO, (C) PNPLA3-KO, (D) PL39-KO, (E) PL38-KO and (F) LPL1-KO. Unpaired students t-test revealed no significant differences between WT and KO parasite lines.

